# The evolution of the temporal program of genome replication

**DOI:** 10.1101/210252

**Authors:** Nicolas Agier, Stéphane Delmas, Qing Zhang, Aubin Fleiss, Yan Jaszczyszyn, Erwin van Dijk, Claude Thermes, Martin Weigt, Marco Cosentino-Lagomarsino, Gilles Fischer

## Abstract

Comparative analyses of temporal programs of genome replication revealed either a nearly complete conservation between closely related species or a comprehensive reprogramming between distantly related species. Therefore, many important questions on the evolutionary remodeling of replication timing programs remain unanswered. To address this issue, we generated genome-wide replication timing profiles for ten yeast species from the genus *Lachancea,* covering a continuous evolutionary range from closely related to more divergent species. The comparative analysis of these profiles revealed that the replication program linearly evolves with increasing evolutionary divergence between these species. We found that the evolution of the timing program mainly results from a high evolutionary turnover rate of the cohort of active replication origins. We detected about one thousand evolutionary events of losses of active replication origins and gains of newborn origins since the species diverged from their last common ancestor about 80 million years ago. We show that the relocation of active replication origins is independent from synteny breakpoints, suggesting that chromosome rearrangements did not drive the evolution of the replication programs. Rather, origin gains and losses are linked both in space, along chromosomes, and in time, along the same branches of the phylogenetic tree. New origins continuously arise with on average low to medium firing efficiencies and increase in efficiency and earliness as they evolutionarily age. Yet, a subset of newborn origins emerges with high firing efficiency and origin losses occur concomitantly to their emergence and preferentially in their direct chromosomal vicinity. These key findings on the evolutionary birth, death and conservation of active replication origins provide the first description of how the temporal program of genome replication has evolved in eukaryotes.

## Introduction

To ensure the completion of genome doubling before cell division, eukaryotic chromosomes initiate DNA replication from multiple sites, termed replication origins. Mammalian and yeast genomes initiate replication at hundreds and thousands of active origins respectively, that are selected from a larger pool of possible Autonomously Replicating Sequences (ARS) ^1–3^. In yeast, active origins are distributed throughout the genome at non-transcribed and nucleosome-depleted sequences and comprise a specific DNA motif called ARS consensus sequence which is bound by the Origin Recognition Complex throughout the cell cycle ^4–6^. Despite of this partially pre-programmed replication activity, different cells in a population may use different subsets of active origins. Indeed, DNA combing and fluorescence microscopy experiments confirmed the stochastic activation of origins at the individual cell level ^7–10^. Nevertheless, at the population level, a specific temporal program of genome replication emerges and can be recapitulated by averaging the heterogeneous replication kinetics of a large number of cells ^7^. Time-course measurements by microarray hybridization or high throughput sequencing of replication progression during S-phase give access, experimentally, to the temporal program of genome replication in a cell population ^11^. Mathematical modeling of such replication timing data gives access to the stochastic firing components of each individual origin ^10,12–14^. First, each replication origin has a given probability of activation, named efficiency, which represents the proportion of the cells in a population in which the origin actively fires. Some origins have high efficiencies, activating in a majority of cells while others have low efficiencies and fire more rarely ^15,16^. Second, each origin has an intrinsic strength called characteristic firing time, which represent its time of activation during S-phase (in the absence of interfering neighboring origins). Some origins fire early, whereas others fire late ^17,18^. Firing time and efficiency both reflect the probability of origin firing, either per unit of time for the former or over the entire S-phase for the latter. Consequently, timing and efficiency are correlated ^16,19,20^, the fundamental difference between the two being the fact that efficiency incorporates the effect of passive replication from forks originated at different origins while firing rate is an origin-specific quantity.

Early microarray-based studies in *Saccharomyces cerevisiae* established the first genome-wide replication program providing replication timing information for all genomic sequences as well as replication origin location along chromosomes ^18,21^. Now, temporal programs of genome replication have been established for many eukaryotic genomes including three closely related *Saccharomyces sensu stricto* species ^22^ and nine more distantly related species including *Candida glabrata, Naumovozyma castellii, Tetrapisispora blattae, Zygosaccharomyces rouxii, Kluyveromyces lactis, Lachancea waltii, Lachancea kluyveri, Pichia Pastoris* and *Candida albicans* ^3,23–27^. However, the question of how the temporal regulation of genome replication has evolved could so far only be tackled in yeast. An early comparative genomics study among closely related *Saccharomyces* species showed that phylogenetic conservation can be used to determine the genome-wide location of replication origins in *S. cerevisiae* ^2^. Other comparative genomic approaches later confirmed the existence of conserved sequence elements necessary for origin function in other yeast genomes as well as a conserved role for centromeres and telomeres in defining early and late origin firing, respectively ^25,26,28–30^. Comparative analysis of replication timing also revealed that at short evolutionary distances, between *Saccharomyces* species, most active origins remained conserved both in location and in activation time, resulting in an important conservation in the temporal order of genome replication ^22^. On the contrary, comparisons between more distantly related species revealed that the conservation of the temporal organization of replication was restricted to specific genetic elements such as centromeres and histone genes that are among the first regions to replicate and telomeres that are among the last ^27^. Only a small proportion of replication origins (5 to 30%) are conserved in position between *S. cerevisiae, L. waltii, L. kluyveri* or *K. lactis* ^3,23,28^. Such a level of reprograming contrasts with the global conservation observed between *Saccharomyces* genomes and precludes any chance to identify the selective forces responsible for the conservation, gain and loss of replication origins over evolutionary time.

To overcome these limitations and to progress in our understanding of the evolutionary remodeling of the temporal programs of genome replication, we focused on the continuous evolutionary range covered by the genus *Lachancea*, ranging from closely related to more diverged yeast species, ^31^. We reasoned that such a continuous range of relatedness should give access to all intermediate evolutionary time points from highly conserved temporal order of genome replication to significantly different cohorts of active replication origins. We first characterized the genome replication dynamics and origin usage at the population level in ten *Lachancea* species. This unique dataset allowed us to infer all events of origin gains and losses since these species diverged from their last common ancestor. We then correlated the functional properties of replication origins from equivalent evolutionary ages, such as their chromosomal location, firing time and efficiency to reveal new rules that govern the birth, death and conservation of active replication origin during evolution.

## Results

### Temporal programs of genome replication in the genus Lachancea

We experimentally measured the temporal programs of genome replication by assessing DNA copy number change during S-phase in the 10 *Lachancea* species with high quality genome assemblies ^31–33^. Cells were synchronized by physical discrimination of G1 cells and synchronously released into S-phase. Time-point samples were taken during S-phase until cells reached the G2 phase and DNA samples were analyzed using Illumina deep sequencing ^34^. The gradual change of DNA copy number was measured to determine the mean replication time, called Trep, along the 10 genomes (**Supplementary Fig. 1, Supplementary Table 1**). To assess the reproducibility of the 10 replication timing profiles we performed a Marker Frequency Analysis (MFA), consisting in measuring replication dynamics directly from an exponentially growing cell population ^35^ and also compared our results to previously published profiles for *L. waltii* and *L. kluyveri* ^3,23^. For each species, the Trep and MFA profiles were highly correlated, indicating a good reproducibility of the experiments (**Supplementary Fig. 2**). In addition, we inferred the location of replication origins along the chromosomes by applying a peak calling method to both the Trep and MFA profiles ^34^ and stringently defined active replication origins only if peaks were co-detected in both profiles (**Supplementary Fig. 2, Supplementary Table 1**).

We identified 2,264 active replication origins in the 10 *Lachancea* genomes (**Supplementary Table 1**). These genomes undoubtedly contain many more dormant origins that have efficiencies too low to be detected by our population sequencing approach. The total number of active origins per genome varies from 200 in *L. cidri* to 256 in *L. fantastica* (**Table 1**). A positive correlation between the numbers of active origins and both the genome and the individual chromosome sizes results in a constant origin density along chromosomes of 1 origin every 47 kb (*R*^*2*^ = 0.83 and *R*^*2*^= 0.95, respectively, **Supplementary Fig. 3a,b**). The distribution of the distances between adjacent origins for all the 10 *Lachancea* genomes revealed that origins are more regularly spaced than what would be expected by chance (**Supplementary Fig. 3c**), as previously described for *S. cerevisiae, K. lactis, L. kluyveri* and *L. waltii* ^36^. We also found common features shared between all 10 replication programs, such as the spatial alternation between early and late replicating large chromosomal regions, with the exception of the left arm of chromosome C in *L. kluyveri* ^23,30^, as well as both early replicating centromeres and histone genes ^27^ and late replicating telomeres (**Supplementary Fig. 1** and **Table 1**).

**Table 1:** Replication profile features

| Species | Genome size | Nb ORI | ORI every ... | Median replication time (min) |  |  | Average S-phase duration (min) |
| --- | --- | --- | --- | --- | --- | --- | --- |
|  |  |  |  | Centromere | Telomere | Histone |  |
| *L. fantastica* | 11.3 Mb | 256 | 44 Kb | 6 +/- 2 | 29 +/- 7 | 3 +/- 3 | 39 |
| *L. meyersii* | 11.3 Mb | 237 | 48 Kb | 2 +/- 2 | 17 +/- 5 | 1 +/- 1 | 24 |
| *L. dasiensis* | 10.7 Mb | 234 | 46 Kb | 4 +/- 2 | 24 +/- 6 | 3 +/- 1 | 36 |
| *L. nothofagi* | 11.3 Mb | 241 | 47 Kb | 6 +/- 4 | 28 +/- 9 | 3 +/- 1 | 42 |
| *L. waltii* | 10.2 Mb | 225 | 46 Kb | 9 +/- 3 | 24 +/- 11 | 7 +/- 1 | 41 |
| *L. thermotolerans* | 10.4 Mb | 211 | 49 Kb | 2 +/- 2 | 16 +/- 5 | 1 +/- 1 | 29 |
| *L. mirantina* | 10.1 Mb | 202 | 50 Kb | 7 +/- 3 | 35 +/- 16 | 1 +/- 1 | 48 |
| *L. fermentati* | 10.3 Mb | 214 | 48 Kb | 2 +/- 4 | 42 +/- 4 | 1 +/- 1 | 47 |
| *L. cidri* | 10.1 Mb | 200 | 51 Kb | 4 +/- 3 | 25 +/- 4 | 2 +/- 1 | 41 |
| *L. kluyveri* | 11.3 Mb | 244 | 46 Kb | 1 +/- 1 | 26 +/- 6 | 1 +/- 1 | 32 |

### A continuous evolutionary range of genome replication timing profiles

In order to determine the degree of conservation of replication profiles between different pairs of species, we calculated the Spearman’s rank correlation coefficients between their replication timing programs. For this purpose, we projected the replication timing of one species onto the chromosomal coordinates of a second species, using synteny conservation between the two genomes (See Methods). We applied this methodology to all pairwise comparisons between the previously published replication datasets in *Saccharomyces* species ^22^, *Candida albicans* ^25^ and our 10 new *Lachancea* replication timing data (**Fig. 1a,b**). We found remarkably high degrees of replication profile correlation between all *sensu stricto Saccharomyces* species, varying from 0.79 to 0.86, as previously reported ^22^. On the contrary, at large phylogenetic distances, all pairwise comparisons involving two species from different genera (*Saccharomyces, Lachancea* or *Candida*) show little to no degree of correlation, preventing any reliable comparative study. Interestingly, pairwise comparisons within the genus *Lachancea* revealed that the Spearman’s rank coefficients stagger from highly conserved replication profiles (*Rho* = 0.74) to more variable programs (*Rho* = 0.41), with all intermediate levels of correlation in-between (**Supplementary Fig. 4**). These coefficients linearly anti-correlate with phylogenetic distances (*R*^*2*^ = 0.88, *P* < 2.2 × 10^−16^), showing that replication timing linearly evolves along with protein divergence (**Fig. 1c**). Such a continuous range of replication programs makes the genus *Lachancea* an ideal candidate to investigate the various causes behind the progressive reprograming of genome replication during evolution.

**Figure 1:**
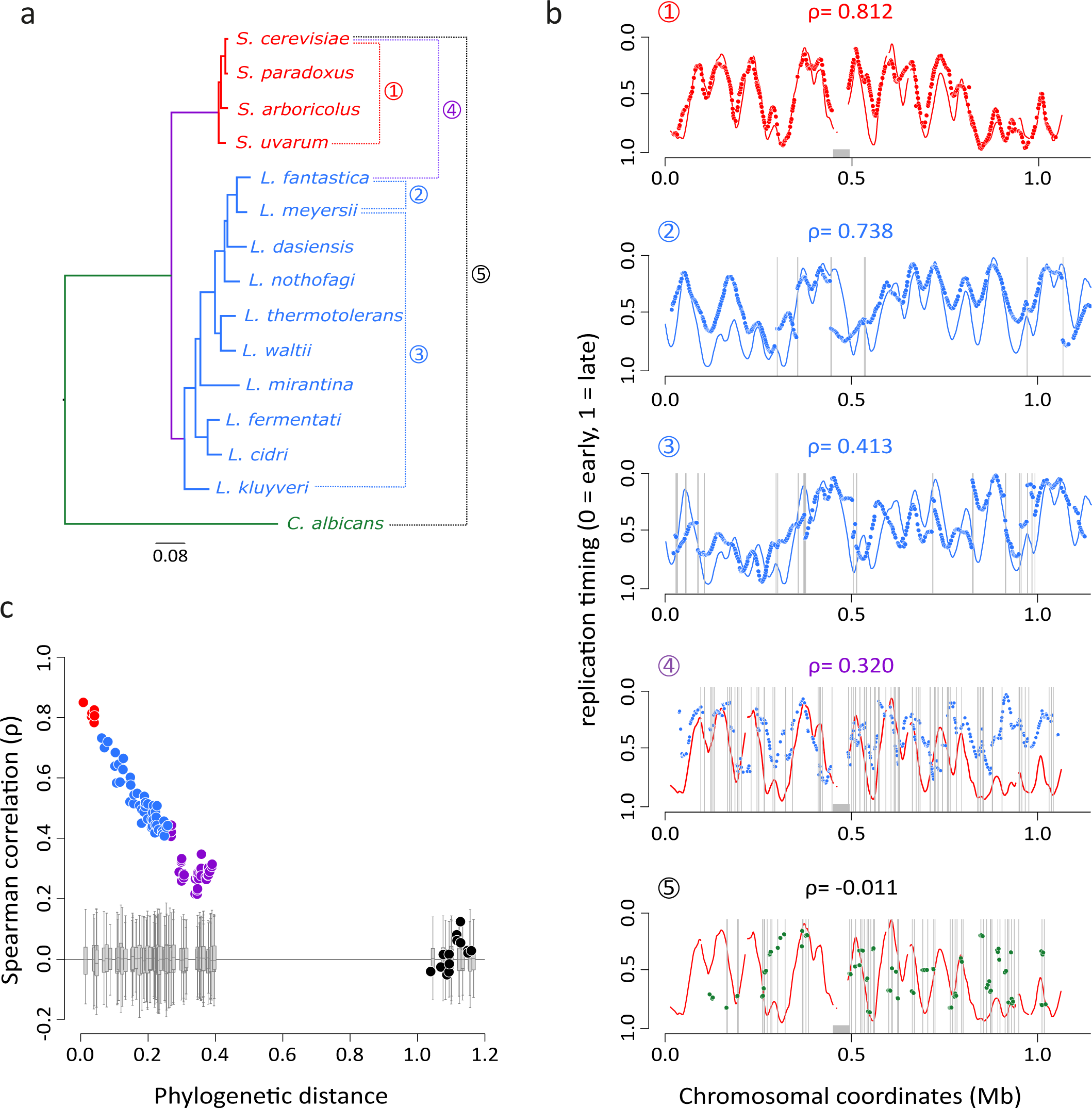
Evolution of replication timing profiles in yeast. **a.** Phylogeny of 15 yeast species inferred from a Maximum Likelihood analysis of a concatenated alignment of 510 protein families. The circled numbers to the right of the tree relate to the pairwise species comparisons presented in the B panel. **b.** Synteny-based projections of replication timing profiles between pairs of species. The color coding is identical to panel A. The Rho value above each profile indicates the genome-wide Spearman’s rank correlation coefficient between the replication timing programs of the two compared species. Vertical grey bars indicate the position of synteny breakpoints between genomes. Comparisons 1, 4 and 5 represent the projections of *S. uvarum, L. fantastica* and *C. albicans* timing data, respectively, onto the replication profile of *S. cerevisiae* chromosome XII. The grey rectangle symbolizes the rDNA locus in this chromosome. The comparisons 2 and 3 represent the projections of *L. fantastica* and *L. kluyveri* timing data, respectively, onto the replication profile of *L. meyersii* chromosome 0D. **c.** Correlation between replication profile conservation and phylogenetic distance. Spearman coefficients on the y-axis correspond to the synteny-based projections of genome replication timing profiles between pairs of species as illustrated in B. Red, blue, purple and black dot series correspond to intra-*Saccharomyces,* intra-*Lachancea, Saccharomyces vs Lachancea* and *C. albicans vs* all other species comparisons, respectively. The grey boxplots show the Spearman correlation values of a null model where, for each pairwise comparison, we applied a random offset to the coordinates of the syntenic genes in one of the two compared genomes. The offset was re-defined 100 times in both directions (offset applied 100 times to both genome 1 and genome 2) and correlations were calculated for each combination of one offset profile from one genome and one original profile from the other genome.

### Differential conservation of active origins drives the evolution of timing programs

We first determined that the number of syntenic homologues, considered as orthologous genes hereafter, that were used to project the replication timing of one genome onto the other was globally constant between all pairs of *Lachancea* species, ruling out the possibility that the anti-correlation we observed in **Fig. 1c** resulted from a bias in synteny detection (**Fig. 2a**). By contrast, the number of synteny blocks clearly increases with phylogenetic distance (**Fig. 2a**) as a result of the accumulation of genome rearrangements during evolution ^31,37,38^. We found that local Spearman’s rank coefficients were on average weaker around synteny breakpoints than within conserved synteny regions, suggesting that genome rearrangements do have a local impact on the evolution of the replication profiles (**Fig. 2b**). Yet, at least part of this decay could be due to a purely technical component because projected profiles at the edge of synteny blocks are necessarily discontinuous. Importantly, the anticorrelation slopes based on the local coefficients both near breakpoints and within conserved synteny blocks are similar to the slope of the global correlation (**Fig. 2b**). This reveals that (i) the local impact of genome rearrangements on replication coefficients is constant over the entire phylogenetic span and (ii) the progressive evolutionary reprograming of the replication profiles is visible within the conserved synteny regions devoid of rearrangements showing that the local effect of genome rearrangements is not sufficient to explain the evolution of the replication programs (**Fig. 2b**).

**Figure 2:**
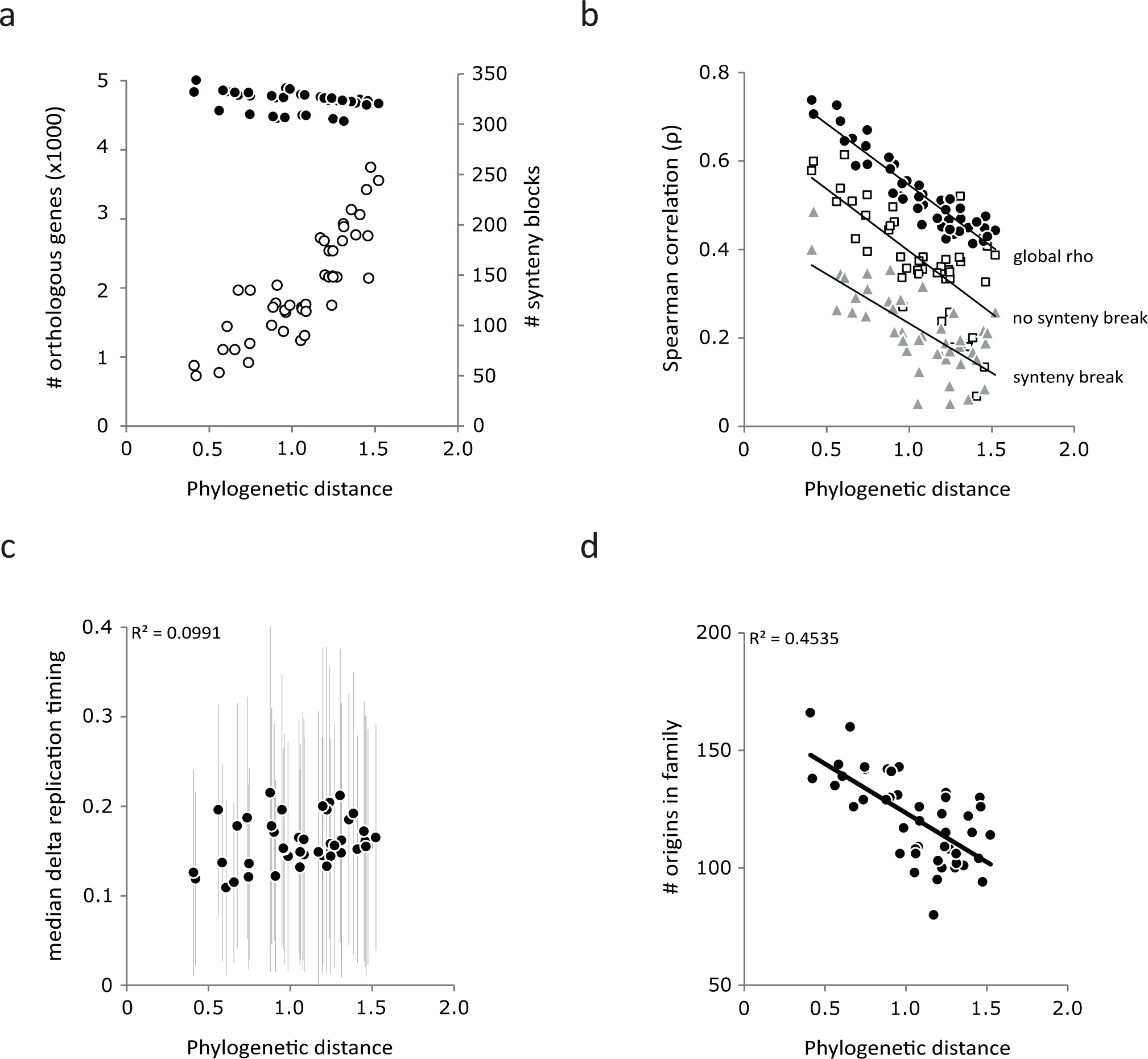
Evolution of genomic properties in the genus *Lachancea*. In all panels, each point corresponds to a pairwise comparison between two *Lachancea* species. The values of the property indicated on the y-axis are plotted as a function of the phylogenetic divergence between all pairs of *Lachancea* species as defined in ^31^. **a.** The number of syntenic homologs and synteny blocks are represented by black and open circles, respectively. **b.** Comparison of global and local correlation coefficients between pairs of genome replication timing profiles. The global coefficient plot (black circles) is identical to that of the blue dots in figure 1C (*R*^*2*^ = 0.868, *P* < 2.2 × 10^−16^). Open squares and grey triangles represent the local correlation coefficients averaged over the same number of regions of 5 genes that are either located 25 genes away from the synteny breakpoints (*R*^*2*^ = 0.502, *P* = 2.24 × 10^−6^) or directly flank the breakpoints (*R*^*2*^ = 0.477, *P* = 1.6 × 10^−6^), respectively. **c.** Origin firing time differences between conserved replication origins in all pairs of species. The black dots indicate the median values for pairwise comparisons and the grey bars show the standard deviations. **d.** Relative conservation of active replication origins in the different pairs of *Lachancea* genomes (*R*^*2*^ = 0.45, *P* = 3.9 × 10^−07^).

We next sought whether the number of replication origins, their differential firing times, or their variable conservation levels could have driven the evolution of the replication timing programs. First, we found no correlation between replication profile conservation and the raw number of active origins in the genomes (**Supplementary Fig. 5**). Second, we constructed families of orthologous replication origins based on synteny conservation (See Methods), to test whether program evolution could have resulted from differential conservation of active replication origins. We excluded 96 subtelomeric origins due to poor synteny conservation and clustered the remaining 2,168 internal origins, representing 96% of the total, into 374 multi-origin families comprising 1,956 origins (90%) and 212 species-specific singleton origins (10%, **Supplementary Fig. 6a**). We found that the differences in origin activity, as measured by peak height on replication timing profiles, between orthologous origins conserved in different species did not correlate with the phylogenetic distances between these species, showing that the reprograming of origin activity had little effect, if any, on the evolution the replication program (**Fig. 2c**). However, we found a negative correlation between the number of conserved replication origins between the pairs of species and their phylogenetic distances (**Fig. 2d**). Given that phylogenetic distances also correlate with replication timing Spearman’s rank coefficients (see **Fig. 1c**), it results that the numbers of conserved origins correlate with the conservation of the replication timing profiles (*R*^*2*^ = 0.57, *P* = 2.5 × 10^−09^). This result shows that the higher the number of conserved orthologous origins between two species is, the more similar their replication programs are. In other words, the appearance and disappearance of active replication origins would be the dominant process for shaping replication profiles during evolution.

### Highly dynamic evolutionary turnover of active replication origins

We reconstructed the evolutionary history of active replication origins along the branches of the phylogenetic tree under a birth/death evolutionary model using *L. kluyveri* as the outgroup species (Methods). We identified 220 origin families, totaling 1,310 origins in the nine *Lachancea* genomes (excluding *L. kluyveri*) that were vertically inherited from the last common ancestor of the clade (designated as *L.A2* in **Fig. 3**). We will refer to them as ancestral origins hereafter. Extant genomes comprise on average 68% of ancestral origins (between 59% in *L. dasiensis* to 83% in *L. fermentati*). Only 37 out of these 220 ancestral families (17%) were faithfully transmitted from *L.A2* without any subsequent loss of origins along the branches of the tree. This means that 83% of the ancestral families underwent at least one event of origin gain or loss, demonstrating that the evolutionary turnover of the cohort of active replication origins is a very dynamic process. Note that what we call here a gain of a new active origin could correspond to the activation of a dormant origin. Similarly, what we call an origin loss could correspond to the inactivation of previously active origin or to the reduction of its activity to a level below the sensitivity of the experiment. We located on the phylogenetic tree a total 916 evolutionary events, including 477 losses of active origins and 439 gains of newborn origins, that modified the repertoire of active replication origins since the nine *Lachancea* species diverged from their last common ancestor (**Fig. 3**). These figures are conservative as they exclude 94 events that occurred on the two most internal branches of the tree, the *b2* and *L. kluyveri* branches, for which it was not possible to discriminate an origin loss on one branch from an origin gain on the other branch. The 439 gain events that occurred along the phylogenetic tree resulted in 623 *Lachancea-*specific origins because a single gain event that occurs on an internal branch of the tree results in several origins present in different descendant species. The 623 *Lachancea-*specific origins represent 32% of all active replication origins present in the 9 *Lachancea* genomes today.

**Figure 3:**
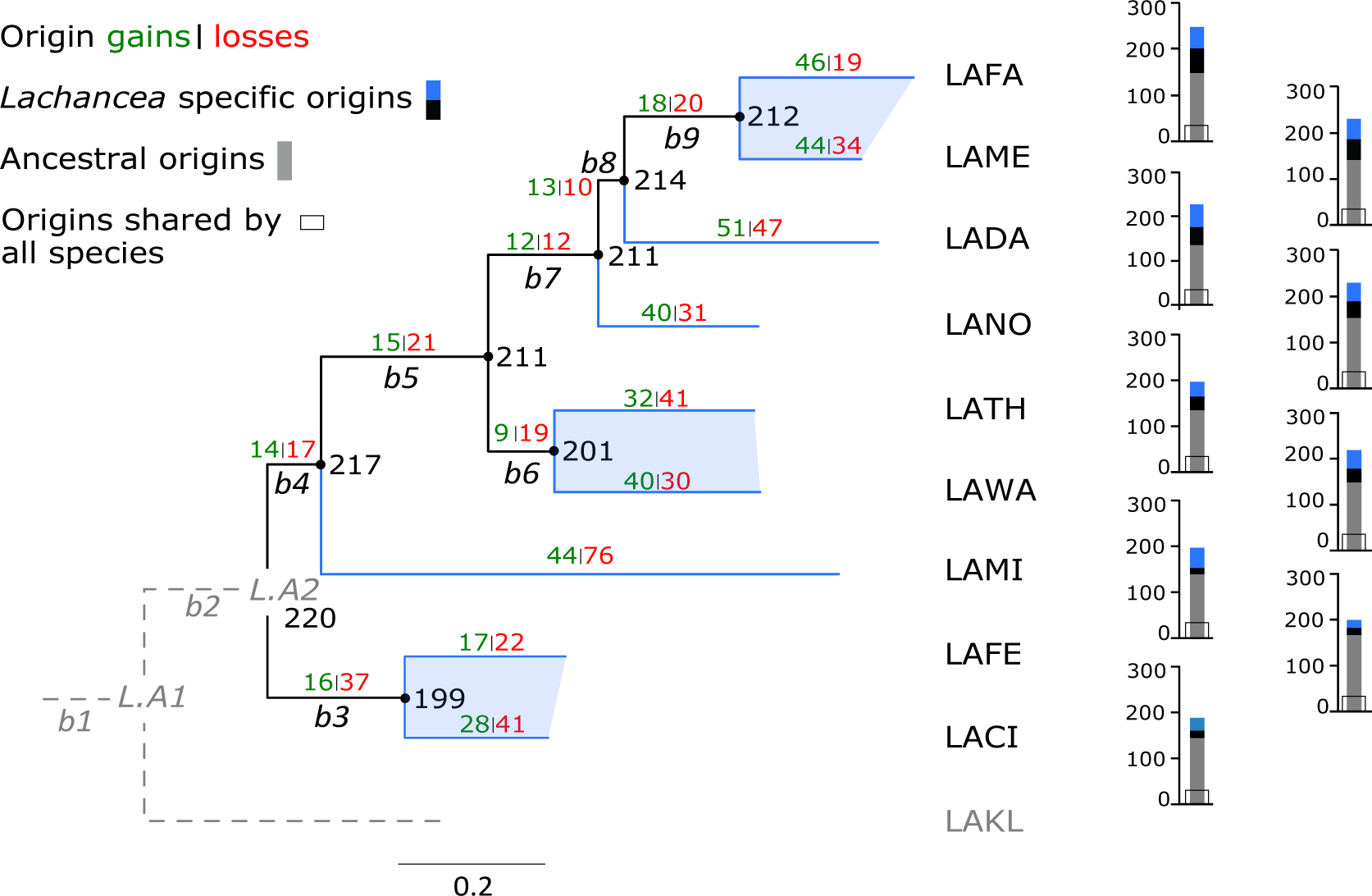
Evolution of the repertoire of active replication origins in the genus *Lachancea.* The phylogeny of the 10 *Lachancea* species is taken from ^31^. Abbreviations of the species names: LAFA = *L. fantastica*, LAME = *L. meyersii*, LADA = *L. dasiensis*, LANO = *L. nothofagi*, LATH = *L. thermotolerans*, LAWA = *L. waltii*, LAMI = *L. mirantina*, LAFE = *L. fermentati*, LACI = *L. cidri* and LAKL = *L. kluyveri*. LAKL was used as the outgroup species, therefore evolutionary events that occurred on both the LAKL and the *b2* branches (dotted lines) could not be retraced. Internal branches, labeled *b3* to *b9,* and terminal branches are drawn in black and blue, respectively. The number of origin gains (in green) and losses (in red) were estimated for each branch of the tree under a Birth-Death evolutionary model (Methods). Histograms on the right indicate the number of active origins that were vertically inherited from the *L.A2* ancestor (in grey), gained on internal branches of the tree (in black) and gained on terminal branches (in blue). The proportion of ancestral origins retained in all nine genomes (not considering LAKL) is indicated by the open frame. The inferred number of active replication origins in the ancestral genomes is indicated next to the corresponding node of the tree.

The number of origin losses per branch, and to a lesser extent the number of gains, significantly correlate with branch length (*R*^*2*^ = 0.78, *P* = 1.1 × 10^−05^ and *R*^*2*^ = 0.37, *P* = 1.5 × 10^−02^, respectively), suggesting that the accumulation of point mutations could have had an impact on the dynamics of the set of active origins (**Supplementary Fig. 7a**). We previously showed that branch lengths also correlate with the number of inversions, translocations and gene duplications, but not with gene losses, in the genus *Lachancea* ^31^. Consequently, origin losses and gains also correlate with inversions, translocations and duplications (**Supplementary Fig. 7b,c**). Although there is no obvious causal relationship between these observations, they reveal that protein divergence, chromosome architecture and replication program evolve in a coordinated manner.

### The loss of an active replication origin depends on its neighboring origin

To further characterize the evolutionary dynamics of active replication origins, we compared the physical and functional properties between conserved, lost and gained replication origins. To make sure that the properties of actual origins reflect as much as possible the properties of the origins at the time they were gained or lost, we focused on the most recent events that occurred along the *Lachancea* phylogeny. Therefore, we concentrated on the six terminal branches that lead to the three most closely related pairs of species, namely *L. fantastica/L. meyersii, L. thermotolerans/L. waltii* and *L. fermentati/L. cidri* (blue shaded area in **Fig. 3**). This dataset comprises 886 origins that have been conserved in at least one pair of sister species, 207 newborn origins that were gained in one genome and 187 previously active origins that were lost in one genome (**Fig. 3**). As physical properties, we studied the distance between each origin and its centromere, its closest telomere, its closest genome rearrangement breakpoints and its closest neighboring origin. As functional properties, we considered the efficiency and the characteristic firing time of each individual origin and that of its closest neighboring origin along the chromosomes. We used a stochastic mathematical model to infer origin firing rates from the fit to our replication timing data and derived the efficiency and firing time of each individual origin in the 10 *Lachancea* genomes from running the model ^14^. Note that for lost origins, we inferred their chromosomal locations, efficiencies and firing times based on the corresponding features of their orthologous origins in the sister genomes where they were retained.

Firstly, we found no difference in the origin-centromere and origin-telomere distances between the conserved, gained and lost origins, showing that their relative chromosomal location did not influence their evolutionary fate (**Supplementary Fig. 8a**). We then tested whether the synteny breakpoints resulting from the 74 rearrangements that occurred along the six terminal branches under consideration ^31^ were associated with the presence of replication origins. We found no clear association between rearrangements and replication origins, neither in the first 5 kb that directly flank the breakpoints nor further away from breakpoints (**Supplementary Fig. 8b**). Similarly, we found no association between origin location and the breakpoints, neither for conserved nor gained nor lost origins (**Supplementary Fig. 8c**). These results both confirm one of our previous findings that replication origins and synteny breakpoints do not significantly collocate in the *L. kluyveri* and *L. waltii* genomes ^23^ and suggest that chromosomal rearrangements play little role, if any, in the evolutionary dynamics of replication origins.

We then estimated the distance separating the conserved, lost and gained origins from their closest neighboring origins or, in the case of lost origins, from the projected position of their orthologous origins. We found that origin losses, and to a lesser extent origin gains, tend to occur much closer to their nearest origin than conserved origins (median distances of 13 kb and 24 kb vs 36 Kb, respectively, **Fig. 4a**). This result clearly shows that origins tend to be lost when they are in close vicinity to another replication origin along the chromosomes. We looked at whether these losses tended to preferentially occur next to conserved, gained or other lost origins by comparing the observed numbers of neighboring origins from the three categories to the expected numbers based on the three sample sizes. We found a strong physical association between origin losses and newly gained neighboring origins (**Fig. 4a right**). Reciprocally, we also found a clear association between newly gained origins and neighboring origin loss positions (**Fig. 4a middle**). These results clearly show that active replication origins are preferentially lost when they are in the close vicinity of a newborn origin that was gained nearby along the chromosome in the same phylogenetic branch of the tree.

**Figure 4:**
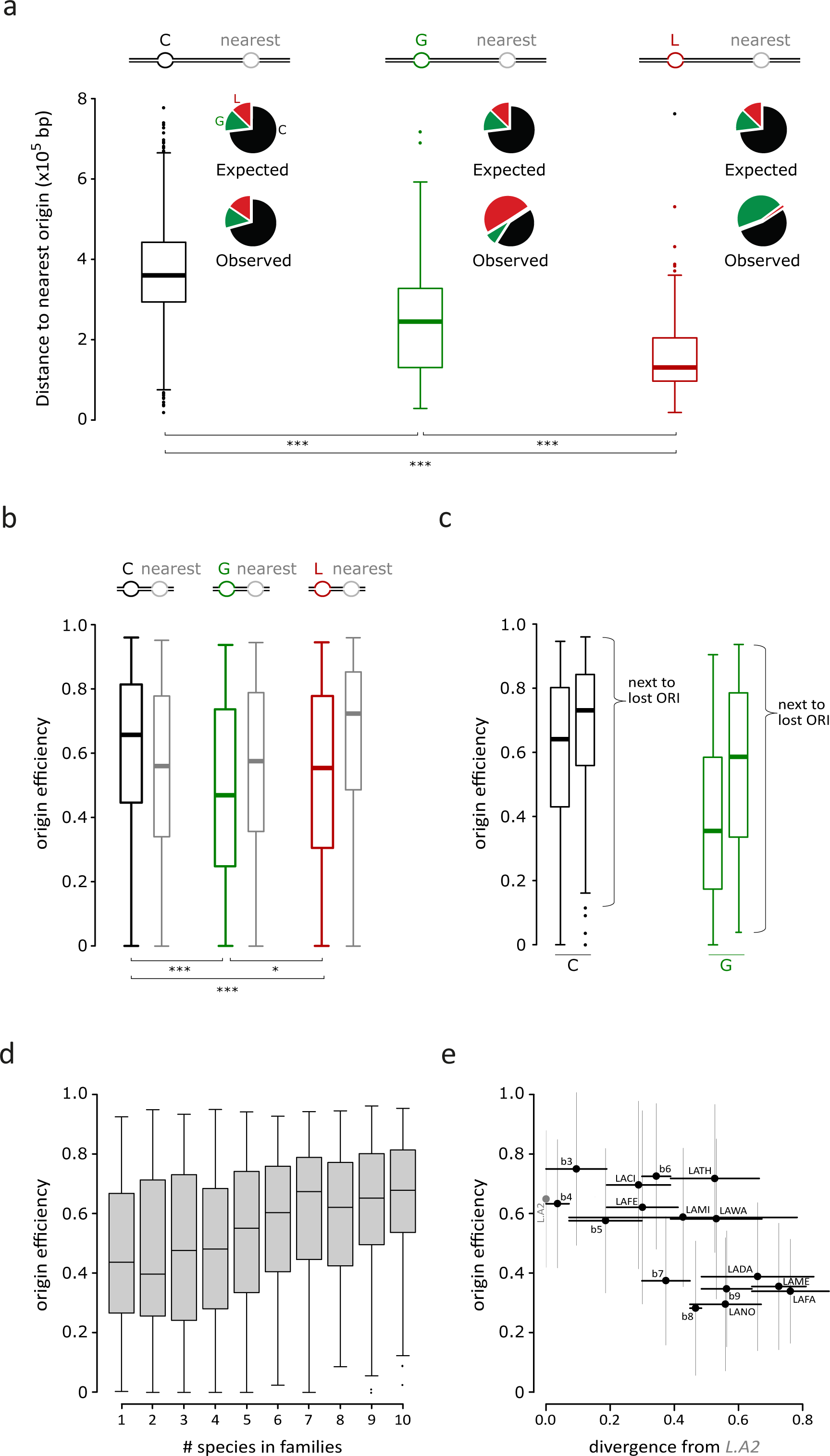
Physical and functional properties of lost, gained and conserved replication origins. In all panels, *** and * represent *P* < 10^−03^ and *P* < 5×10^−02^, respectively, using a Chi-Square Two Sample Test. **a.** Distribution of the distance separating Conserved (C), Gain (G) and Lost (L) origins from their nearest origins. The pie charts show the expected (i.e. the theoretical percentages if the nearest origins were randomly sampled in the population of origins) and the observed proportions of C, G and L nearest origins for the three categories. **b.** Distribution of the efficiencies of C, G and L origins and of their nearest origins. **c.** Split distributions of the efficiencies of conserved and gained origins based on the category of their nearest origin. **d.** Distribution of the firing efficiencies as a function of the number of *Lachancea* species comprised into the families of orthologous origins (*R*^*2*^ = 0.891 on median values, *P* = 4.1 × 10^−05^). **e.** Correlation between origin efficiency and origin age for 546 origin families comprising 220 ancestral and 326 *Lachancea-*specific families (*R*^*2*^ = 0.44, *P* = 1.3 × 10^−03^). Each dot represents the median efficiency of all the replication origins that appeared in a given branch of the phylogenetic tree. The *x*-axis represents the total branch length between *L.A2* (the last common ancestor of the 9 species, indicated in light grey) and the branch of appearance of the new origins, placing the most ancestral origins on the left and the youngest on the right of the plot. Vertical error bars represent the standard deviation of origin efficiencies and horizontal bars represent the span of the branch lengths based on the phylogenetic tree in Figure 3.

Next we compared the efficiencies between the three categories of origins and found that gained and to a lesser extent lost origins have significantly lower efficiencies than conserved origins (**Fig. 4b**). We also found that the efficiencies of the nearest origins that directly flank the lost origins are significantly higher than those flanking both gained and conserved origins. At first sight this result may seem paradoxical because it suggests that an origin would be preferentially lost when it is flanked by a gained and highly efficient origin, while gained origins have in general the lowest efficiencies of all three categories (**Fig. 4b**). However, we found that the subset of gained origins that are in close vicinity to the lost origins include the most efficient of all newborn origins (**Fig. 4c**). The same is true for conserved origins, the ones that are associated with an origin loss being the most efficient of all conserved origins (**Fig. 4c**).

To estimate to which extent the proximity of a highly active origin may have hidden a lower efficiency origin that would have been readily detectable if it was more isolated, we looked for the presence of a replication origin at the location of our predicted origin losses in an independent origin dataset previously generated in *L. waltii* ^3^. In this work, the authors reported the identification of 194 ARS, 156 of which were shown to robustly function as chromosomal replication origins during S-phase. We found that only 5 out of the 30 origin losses that we identified in the *L. waltii* branch (**Fig. 3**) did correspond to a chromosomally active origin in the Di Rienzi and collaborators dataset (17%). In comparison, 120 out of the 156 active origins were also detected as replication origins in our experiments (77%). Therefore, even if a minority of our loss events could actually correspond to low efficiency origins that escaped detection due to the proximity of a highly active origin, the vast majority of the detected events probably correspond to true origin losses.

Therefore, a clear picture of the evolutionary dynamics of replication origins emerges from these analyses, where origins are preferentially lost when located close to a newborn origin that emerged with a high firing efficiency. Similar results were obtained with characteristic firing times, as late firing origins are preferentially lost near early firing origins that have emerged recently on the same phylogenetic branch (**Supplementary Fig. 9**)

### The efficiency and firing time of active replication origins depend on their evolutionary age

We found that efficiencies positively correlate with the number of species that are represented in the origin families (**Fig. 4d**), suggesting that efficiency would be related to the evolutionary age of the origins. We checked that all individual lineages contributed to this signal, as the efficiency of species-specific origins in each species is significantly lower than that of ubiquitous origins that are conserved in the 10 species (**Supplementary Fig. 10a**). We also checked that this relationship between efficiency and origin family size did not result from a lack of precision in origin location for weak and isolated origins as compared to strong and ubiquitous origins (**Supplementary Fig. 10c,d**). However, we reasoned that the number of species per family was not necessarily the best proxy for the evolutionary age of the origins. For instance, a family that comprises 4 species could nevertheless be older than a family that comprises 5 species if the former underwent more origin losses on different branches of the tree than the latter. In addition, species-specific origins could in fact have very different ages because of the very large length variations between the terminal branches of the tree. In order to remove these putative confounding factors, we used our reconstruction of replication origin history (**Fig. 3**) to assign a phylogenetic age to each origin family, corresponding to the cumulated branch lengths between *L.A2* and its branch of origination in the phylogenetic tree. We found a significant correlation between the phylogenetic age and the efficiency of the origin families (**Fig. 4e**). Both ancestral and oldest *Lachancea-*specific origins, originating from *L.A2* and the *b3/b4* branches respectively, fire more efficiently on average than the youngest origins that were gained on the small terminal branches leading to *L fantastica* and *L. meyersii* (**Fig. 4e**). The corresponding analyses performed with characteristic firing times gave consistent results with older origins firing on average earlier than younger origins (**Supplementary Fig. 11**). Therefore these results show that origin activity correlates with the evolutionary age of the replication origins.

## Discussion

The present study is the first detailed reconstruction of the evolutionary history of genome replication in eukaryotes. By focusing on the model yeast genus *Lachancea,* which exhibits a continuous evolutionary range, from closely related species to more divergent genomes, we captured all intermediate states between highly conserved and significantly reprogrammed temporal orders of genome replication. The replication-timing program evolves in a coordinated manner with protein sequence and chromosome architecture (**Fig. 1c** and **Supplementary Fig. 7**). However, we discovered that surprisingly the accumulation of chromosomal rearrangements did not drive the evolution of the replication program (**Fig. 2b** and **Supplementary Fig. 8b,c**). Rather, we found that the dominant process in replication profile evolution is the appearance and disappearance of chromosomally active replication origins, leading to a highly dynamic evolutionary renewal of the cohort of replication origins across species (**Fig. 2d**). The evolutionary dynamics of replication origins in *Lachancea* far exceeds that of chromosome rearrangements with 394 recent events of origin gains and losses compared to only 74 translocations and inversions that reached fixation on the terminal branches of the tree ^31^(**Fig. 3**). The small proportion, 17%, of ancestral origin families that remained conserved across all *Lachancea* genomes also illustrates the high evolutionary turnover of active replication origins. These results are in marked contrast with the high conservation in the location and activation time of chromosomally active origins between much more closely related species from the *Saccharomyces* sensu stricto complex ^22^. In total, we characterized 1,010 origin gains and losses since the *Lachancea* species diverged from their last common ancestor.

The chromosomally active replication origins detected by our genome-wide timing survey only correspond to a small subset of a larger pool of possible ARS because cells, in order to overcome potentially irreversible double fork stalling events, license many more origins than they normally use ^1–3,39–41^. For instance, in *S. cerevisiae*, the experimental deletion of all efficient replication origins from entire chromosomes only causes a marginal mitotic instability because dormant origins that are normally chromosomally inactive become active and contribute to the replication of the chromosome ^42,43^. The evolutionary dynamics of replication origins uncovered in this study is not directly comparable to this ARS modularity because time scales are very different. However, we cannot rule out that the evolutionary gains of new chromosomally active origins could in fact correspond to the activation of dormant ARS rather than true emergence of new origins.

The key result of our study is that we uncovered several principles that governed the gain, loss and conservation of replication origins during genome evolution. Firstly, new chromosomally active replication origins are continuously gained and lost during genome evolution (**Fig. 3** and **Supplementary Fig. 7**). Secondly, the activity of replication origins depends on their evolutionary age. New origins that were gained on terminal branches of the tree emerge with globally low efficiencies and late firing times (**Fig. 4b** and **Supplementary Fig. 9a**), conserved origins that appeared on internal branches have intermediate activity and ancestral origins are the strongest of all origins (**Fig. 4e** and **Supplementary Fig. 11b**). These findings imply that present-day genomes comprise origins of various strengths depending on their evolutionary age. They also suggest that origin activity becomes stronger while they age over long evolutionary periods. An alternative possibility would be that natural selection would rapidly purge the genomes from low efficiency replication origins and select for the few new origins that emerged with high efficiency. However, the latter hypothesis seems less likely than the former one because of the following observations. New origins are gained with relatively low activity however, among them, the subset corresponding to the most efficient and earliest origins is physically associated with the lost origins, both in terms of physical distance along the chromosome and time of appearance in the same branch of the phylogenetic tree (**Fig. 4a,c** and **Supplementary Fig. 9b**). Similarly, conserved origins that flank lost origins are also on average the most efficient of all conserved origins (**Fig. 4c**). Reciprocally, origin losses preferentially occur when they are in close vicinity to strong origins (**Fig. 4a,b**). In addition, lost origins have on average intermediate activity as compared to conserved and gained origins but they cover the entire range of efficiency, from weak to strong origins (**Fig. 4b**).

Altogether, the above observations fit within a two-pathway model of coordination of origin gains and losses during evolution that we call loss-first and gain-first. In the loss-first pathway, the main cause explaining the gain of a strong chromosomally active origin would be the initial loss, nearby along the chromosome, of a strong origin, generating a large region devoid of initiation zone (**Fig. 5**). Conversely, in the gain-first pathway, the main cause explaining the evolutionary loss of a weaker replication origin would be the appearance, nearby along the chromosome, of a strong replication origin (**Fig. 5**). In a nutshell, all the above observations fit into a model of evolution of the temporal program of genome replication that relies on four simple principles (**Fig. 5**): (i) chromosomally active replication origins are continuously gained and lost during evolution; (ii) newly gained origins have on average a low activity and their strength increases over evolutionary times; (iii) origin losses are tightly linked to origin gains both in space, along chromosomes, and in time, along the same branches of the phylogenetic tree and (iv) both newly gained and conserved origins that directly flank the origin losses are the most active replication origins.

**Figure 5:**
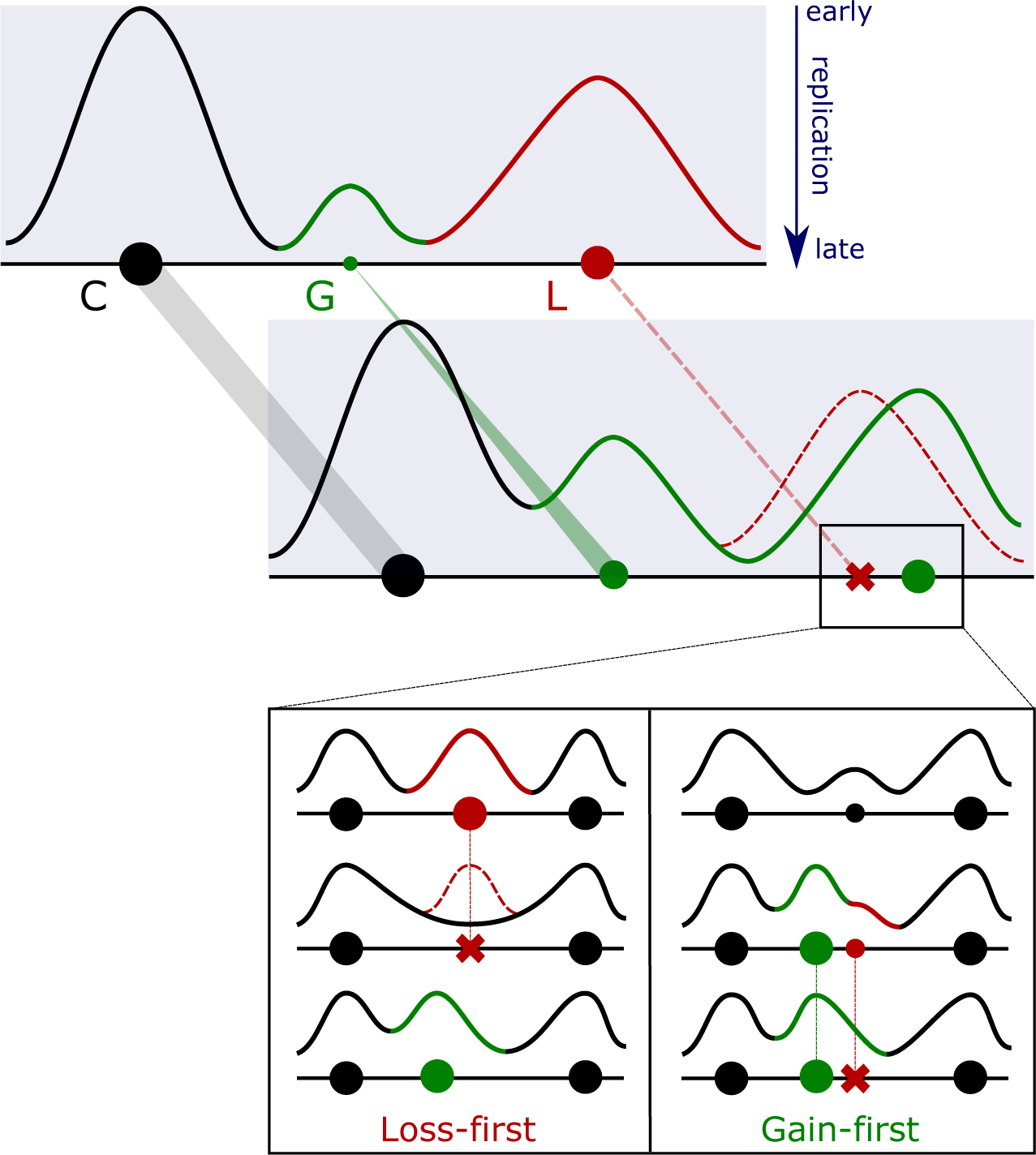
Evolutionary model of the temporal program of genome replication. The diagram on the top illustrates that (i) conserved origins (C) are more ancestral and have on average stronger activity than younger origins, (ii) chromosomally active replication origins are continuously gained (G) and lost (L) during evolution and (iii) newly gained origins have on average low activity and their strength increases over evolutionary times. The two pathways at the bottom, called Loss-first and Gain-first, describe at shorter evolutionary distances the evolutionarily coordinated gain of a strong origin and loss of a strong (left) or weak (right) origin.

It is intriguing to speculate on the physiological principles and constraints leading to the evolutionary tradeoffs that lead to these specific “rules”. It is noteworthy that despite their high evolutionary turnover, chromosomally active replication origins remain more regularly spaced than expected at random in all ten *Lachancea* genomes since they diverged from their last common ancestor approximately 80 million years ago (**Supplementary Fig. 3c**). Regular spacing of replication origins along chromosomes was also reported for other yeast species, including *S. cerevisiae* and *Schizosaccharomyces pombe* ^36,44–46^. These findings suggest that origin distribution has been optimized to limit large inter-origin distances where irreversible double fork stall events are more likely to occur ^36^. If natural selection acts to keep a set of active replication origins regularly spaced along chromosomes, it means that there are costs for keeping origins too close, as well as for keeping a higher and lower density of active origins. However, the molecular determinants that drive the evolutionary loss and gain of active replication origins and therefore the remodeling of the temporal program of genome replication remain so far unidentified.

## Methods

### Yeast strains and growth conditions

All yeast strains and growth conditions are summarized in Supplementary table 1. For all *Lachancea* species, excluding *L. nothofagi* and *L. fantastica*, cells were grown at 30°C in YPD broth (BD-Difco). *L. nothofagi* and *L. fantastica* were grown at 24°C since the former does not grow and the latter aggregates at 30°C. All time-course experiments were performed at 23°C in YPD.

### DNA sample preparation

The experimental and analytical methods used for the time course experiments are fully described in ^34^. Briefly, G1 cells were isolated from an asynchronous cell culture using centrifugal elutriation, and then grown at 23°C in YPD. Time-point samples were taken regularly until the cells reached the G2 phase. Progression of the cells through the cell cycle was monitored using flow cytometry, as described in ^23^. Samples covering the whole S-phase were selected and DNA was extracted using the genomic-tip 20/G isolation kit (Qiagen).

For the expos/stat experiments, cells were grown in YPD at the appropriate temperature. The exponential (Expo) and stationary phase (Stat) samples were collected after 4 and 30 hours of growth. DNA was extracted using the genomic-tip 20/G isolation kit (Qiagen).

### Deep sequencing

For each sample, a minimum of 300ng of genomic DNA was sequenced as 50 bp single reads using Illumina technology (Single-End, 50 bp). A minimum of 10 million and 15 million reads per sample were used for the time course experiment and the Expo/Stat experiment, respectively. To avoid differential PCR biases between samples in the time course experiment, all multiplexed libraries were pooled before PCR amplification as described in ^23^. Libraries were de-multiplexed and adaptator sequences were removed from the reads. Sequences were remapped to the reference genomes ^31–33^ using BWA (0.59) and allowing no mismatch and no gap. Mapped reads were subsequently filtered to keep only unique match and high quality mapping scores (MAPQ > 37, i.e., base call accuracy > 99.98%).

### Mean Replication Time and MFA Profiles

Mean replication times were calculated from time course experiment sequences. Reads were counted in 500 bp non-overlapping windows, along the genome. Changes in DNA copy number were measured by calculating the ratio of the number of sequences between S-phase samples and the reference sample, corresponding to G1 or G2 phase. For each time point, the median of the S/G1 or S/G2 ratio was adjusted to correspond to the DNA content measured by flow cytometry during S-phase progression. Subsequently, ratios were re-scaled between 1 and 2 and for each window, the time where the scaled ratio equaled 1.5 was defined as the Trep. Finally, mean replication times were obtained by smoothing the data with a loess regression (see ^34^ for details).

Marker Frequency Analysis (MFA) profiles were calculated from the Expo/Stat experiments. Reads were counted in all 500 bp non-overlapping windows, along the genome. Windows where the number of sequences was defined as an outlier (> or < 1.5 times interquartile spaces) were filtered out. The MFA ratio is calculated by dividing, for each window, the number of sequences from the Expo sample by the number of sequences from the Stat sample as described in ^35^. The MFA profile is obtained by smoothing the data with a loess regression. All timing data are available in the **Supplementary Table 1**.

### Identification of Replication Origins

The mean replication times and MFA profiles were plotted as a function of the chromosomal coordinates. The first derivative of these curves was estimated by calculating the slope of each coordinate *x* of the window *i* along the genome, using the following formula (*y_i_ − y_i-1_*)/(*x_i_ − x_i-1_*), as described in ^34^. The second derivative was then estimated from the first derivative values using the same method and plotted as a function of the chromosomal coordinates. Both peaks and shoulders from the original mean replication time and MFA curves appear as clear peaks on the second derivative plot. The chromosomal location of these peaks was determined where the slope curves corresponding to the third derivatives intersect 0 from negative to positive value as described in ^34^. We defined peak coordinates as the location of active replication origins only when peaks were co-detected in both the mean replication and the MFA second derivatives profiles. Then, for each replication origin location, we compared its two chromosomal coordinates issued from the two curves to estimate the precision of our experiments in calling origin locations. We found a median and a mean difference between the two peak coordinates of 4.0 kb and 4.9 kb, respectively. All origin locations are available in the **Supplementary Table 1**.

### Comparison of temporal programs of genome replication

The identification of orthologous genes and the construction of synteny blocks were computed with the SynChro algorithm ^47^ for all pairwise combinations between the 10 *Lachancea* species, the 4 *Saccharomyces sensu stricto* species and *C. albicans*. Genome annotations were downloaded from GRYC (http://gryc.inra.fr), saccharomycessensostricto (http://www.saccharomycessensustricto.org) and CGD (http://www.candidagenome.org). We used our mean replication timing profiles for the *Lachancea* species and the previously published replication data for the *Saccharomyces* species and *C. albicans* ^22,25^. For each pairwise combination, a Spearman’s correlation coefficient (rho) is calculated on replication timing for all orthologous pairs. As a null model, the same correlation was calculated after applying an offset on orthologous coordinates for one of the two species of the comparison. The offset was re-defined 100 times and correlations were calculated for each combination of an offset profile from one species and an original profile from the other species.

### Construction of families of conserved orthologous replication origins

We used synteny conservation between the 2 protein coding genes that flank each origin position to construct families of orthologous active origins. Because of the poor synteny conservation in subtelomeres, we excluded the 96 subtelomeric origins that were interstitially located between the telomeres and the first synteny blocks comprising at least 5 syntenic genes. We constructed origin families for the remaining 2,168 internal origins, representing 96% of the total number of origins (**Supplementary Fig. 6a**). For all pairwise comparisons between two *Lachancea* species, we first projected the position of each replication origin of one genome onto the chromosomal coordinates of a second genome based on the synteny conservation of its two flanking coding genes, and reciprocally from the second genome to the first one. Each projection was associated with its nearest resident origin and then, two origins were defined as conserved between two species when the projected and the resident origins were located at most two syntenic genes apart, in both directions (delta=2, see below and **Supplementary Fig. 6b**). The resulting 3,730 pairs of conserved orthologous origins were subsequently clustered into origin families by transitivity. Active replication origins distributed into 374 multi-origin families comprising 1,956 origins (90%) and 212 species-specific singleton origins (10%, **Supplementary Fig. 6a**).

To assess the quality of our origin families, we generated a null model where the positions of the origins were randomized, according to the following constraints:

- randomized inter-origin distances must preserve the same distribution than the observed inter-origin distances. This property is particularly important, since actual origins are more regularly spaced than randomly distributed origins (**Supplementary Fig. 3c**).
- the position of the first origin on each chromosome must be located between the beginning of the chromosome and two-times the position of the first actual origin.
- the relative proportions of intra- and inter-genic origins must be conserved by the randomization procedure.

In particular the first two constraints guarantee that the average number of origins is globally conserved, even if it may slightly fluctuate between samples drawn randomly from the null model.

Thereafter, for each pairwise comparison, origin positions were randomized 100 times in the first genome and projected on the second genome, and reciprocally. Randomized origin families were then constructed with the same procedure as for the real origin families (see above). We used the randomized families to define the optimal number of syntenic genes allowed between two conserved origins. The threshold of delta=2 syntenic genes was determined after testing all values between 0 and 6 intervening genes and looking for the value that maximized the differences between real origin families and null model. We found that Delta=2 was the value that jointly (i) maximized the number of conserved origins in the real versus the random dataset, (ii) maximized the number of families comprising 10 origins (one per species) in real versus random dataset comparisons and (iii) minimized the number of families with more than 10 members (**Supplementary Fig. 6b**). Finally, we also found that Delta=2 limited the proportion of families with more than one origin per species (1.9% for Delta=2 *vs* 8% for Delta=3).

To check that our methodology indeed captured a true evolutionary signal corresponding to the orthology relationship between replication origins, we compared the distributions of the number of origins per family between the real and the random dataset and found that they were clearly different from what is expected from a null model (**Supplementary Fig. 6c**). Moreover, we constructed two phylogenetic trees based on the composition of the real origin families in the 10 *Lachancea* species. First, we performed hierarchical clustering implemented in MEV (http://www.tm4.org./#/welcome) based on the phylostratigraphic patterns of origin families. Second, we built a distance matrix representing the proportion of conserved origins between any two pairs of species and generated a NJ tree, using Phylip version 3.695 ^48^. The two resulting tree topologies are very similar to the topology of the reference species tree based on the concatenation of 3,598 orthologous protein sequences ^31^, with only few bipartitions being different (**Supplementary Fig. 6d**).

### Inference of replication origin history

Based on the composition of the origin families, the evolutionary history origin conservation, gain and loss was inferred using the Gloome online program ^49^. As input, we used simplified phyletic patterns limited to the presence (1) or absence (0) of an origin family within *Lachancea* species. We used an evolutionary model where the probability of gain and loss is equal across all sites. We tolerated more than one possible creation event per family. The parsimony cost of the gains was set to 2. Other parameters were set to default values.

### Data acces

The fastq files are deposited in the Sequence Read Archive under the projet number SRP111158. The following link will direct reviewers to the metadata, including submission structure and sample metadata: ftp://ftp-trace.ncbi.nlm.nih.gov/sra/review/SRP111158_20171010_154743_3d522deaf85577451c01974654b36ad3

## Acknowledgements

This work was supported by the Agence Nationale de la Recherche (GB-3G, ANR-10-BLAN-1606 and Phenovar, ANR-16-CE12-0019). We thank our colleagues, Gianni Liti, Bertrand Llorente, Carolin Müller, Conrad Nieduszynski and Samuel O’Donnell for fruitful discussions and constructive suggestions.

